# Boreal marine fauna from the Barents Sea disperse to Arctic Northeast Greenland

**DOI:** 10.1101/394346

**Authors:** Adam J. Andrews, Jørgen S. Christiansen, Shripathi Bhat, Arve Lynghammar, Jon-Ivar Westgaard, Christophe Pampoulie, Kim Præbel

## Abstract

As a result of ocean warming, the species composition of the Arctic seas has begun to shift in a boreal direction. One ecosystem prone to fauna shifts is the Northeast Greenland shelf. The dispersal route taken by boreal fauna to this area is, however, not known. This knowledge is essential to predict to what extent boreal biota will colonise Arctic habitats. Using population genetics, we show that Atlantic cod (*Gadus morhua*), beaked redfish (*Sebastes mentella*), and deep-sea shrimp (*Pandalus borealis*) specimens recently found on the Northeast Greenland shelf originate from the Barents Sea, and suggest that pelagic offspring were dispersed via advection across the Fram Strait. Our results indicate that boreal invasions of Arctic habitats can be driven by advection, and that the fauna of the Barents Sea can project into adjacent habitats with the potential to colonise putatively isolated Arctic ecosystems such as Northeast Greenland.

## Introduction

The Arctic is warming more rapidly than any other geographical region^1^. Increases in water temperature and loss of sea-ice^2,3^ are expected to induce a northward range expansion of boreal fauna^4–6^, a phenomenon that is already apparent in the Barents Sea^7–9^. Atlantic mackerel (*Scomber scombrus*) exemplifies this trend, having recently displayed an exceptional northward shift in distribution to Spitsbergen^10^. Such predatory newcomers and competitors pose a genuine threat to native Arctic fauna and thus to Arctic ecosystems as they may restructure trophic relationships if their occurrence becomes persistent^8,11,12^.

In 2015 and 2017, boreal species, i.e. juvenile Atlantic cod (*Gadus morhua*), juvenile beaked redfish (*Sebastes mentella*), and adult deep-sea shrimp (*Pandalus borealis*), were observed on the Northeast (NE) Greenland shelf (latitudes 74-77 °N, Fig. 1) for the first time since surveying began in 2002^13^. This was well outside of their known distribution ranges^4,14,15^ (Fig. 2a,d,g). However, the route by which the three species had reached NE Greenland was unknown. The present study aims to determine their population of origin using genetic markers. This knowledge will allow us to infer the dispersal routes taken by these species – information that is critical to forecast (e.g. ^5^) which boreal species may disperse into the Arctic and to what extent they may colonise Arctic habitats^16^.

**Fig. 1.**
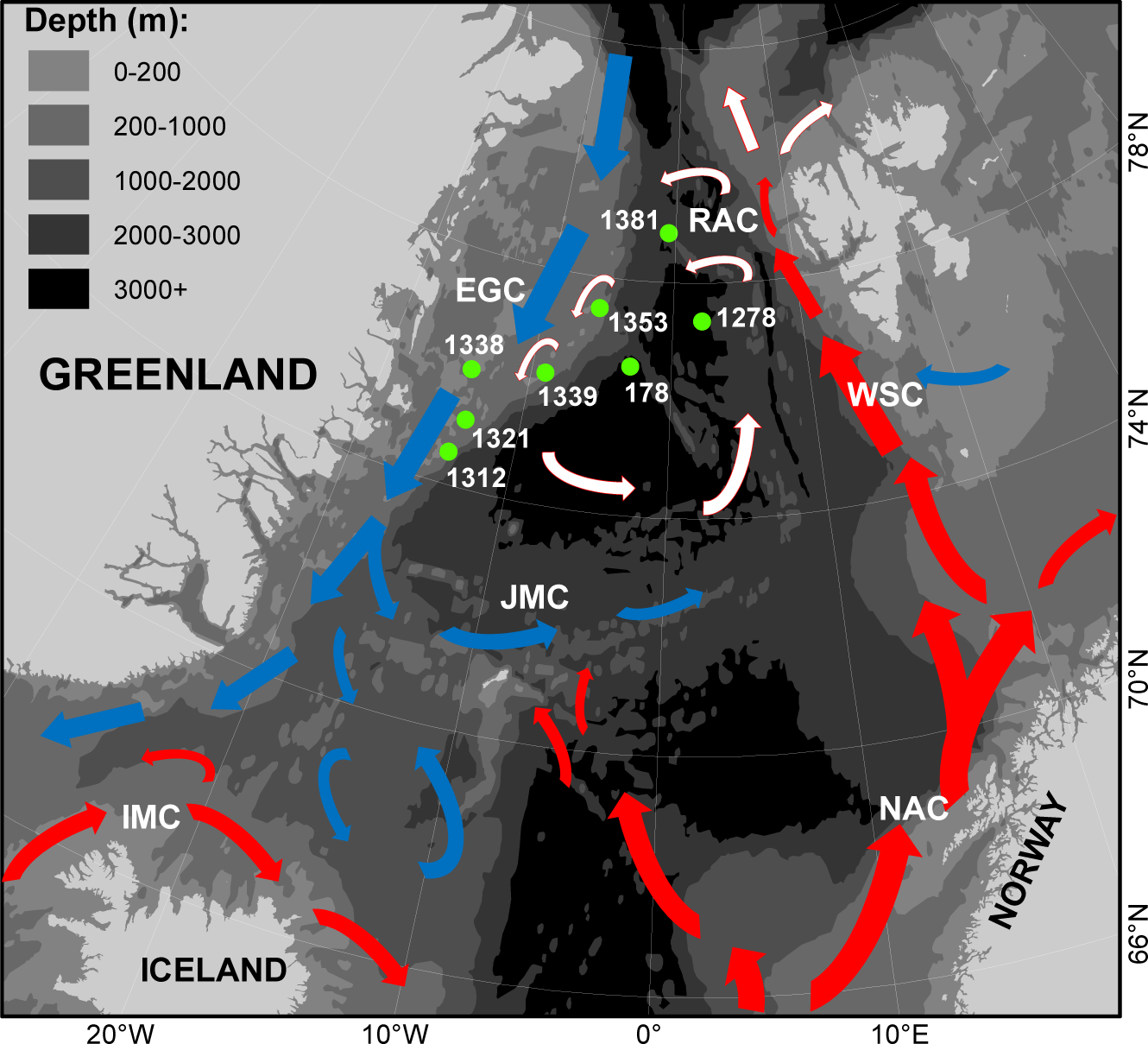
Stations (green full circles) of observation for Atlantic cod (*Gadus morhua*), beaked redfish (*Sebastes mentella*) and deep-sea shrimp (*Pandalus borealis*) (Methods, Table 1). Arrows indicate ocean currents (Source^17,24^). Atlantic surface currents (red arrows): IMC (Irminger Current), NAC (Norwegian Atlantic Current), WSC (West Spitsbergen Current), RAC (Return Atlantic Current). Atlantic sub-surface water (white arrows). Arctic surface currents (blue arrows): EGC (East Greenland Current), JMC (Jan Mayen Current). Arrow size indicates velocity. Map created using ESRI ArcMap (v. 10.6, www.arcgis.com).

**Fig. 2.**
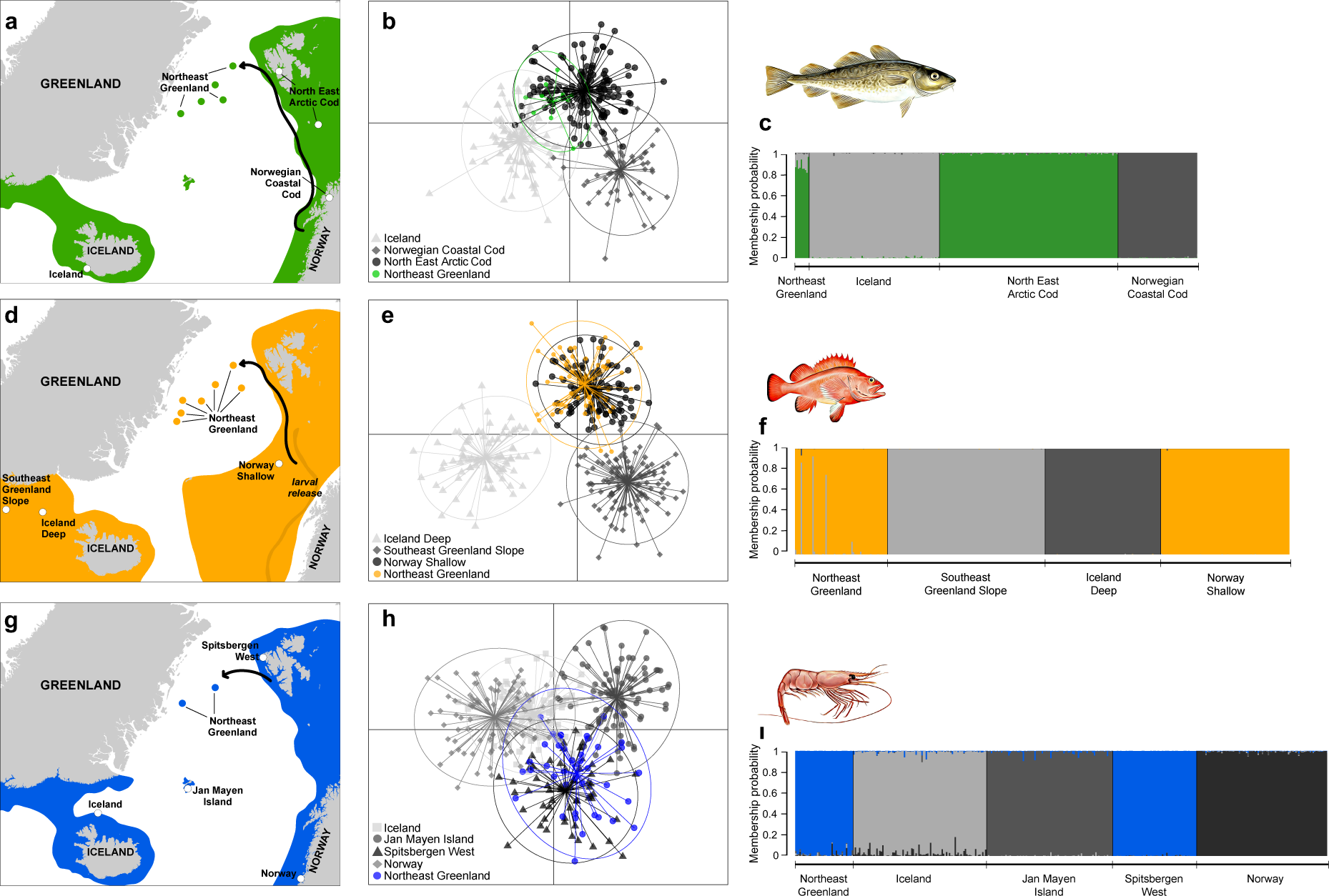
Genetic evidence of Atlantic cod (*Gadus morhua)* (**a, b, c**), beaked redfish (*Sebastes mentella)* (**d, e, f**) and deep-sea shrimp (*Pandalus borealis)* (**g, h, i**) specimens found off Northeast Greenland originating from the Barents Sea. Maps (**a, d, g**) show species known distribution extent (shaded colours) in the Northeast Atlantic, catch sites of individuals in Northeast Greenland (NEG) waters (full circles), reference samples (hollow circles) and a proposed dispersal route (arrow). DAPC scatterplots (**b, e, h**) show how the NEG groups relate to the reference populations of the Northeast Atlantic Ocean. DAPC cluster ellipses were set to contain 95% of genotypes. DAPC scatterplots explain 94% (**b**), 92% (**e**) and 97% (**h**) of the total variation observed. STRUCTURE barplots (**c, f, i**) show membership probabilities (*q*) for NEG individuals based on the reference populations used. Maps were created using ESRI ArcMap (v. 10.6, www.arcgis.com).

We consider two main routes of dispersal to NE Greenland, either 1) via migration against the East Greenland Current^17^ from Iceland, or 2) via advection^18^ from the Barents Sea by the westbound Return Atlantic Current^17,19,20^ (Fig. 1) across the abyssal plains of the Fram Strait. The Norwegian Atlantic Current, along the Norwegian coast, and the West Spitsbergen Current^17,20^, along the Barents Sea shelf-break, are already known to result in the advection of cod, redfish, and shrimp offspring from the Norwegian coast and the Barents Sea proper to Spitsbergen, east of the Fram Strait^21–23^.

## Results

We found that all cod (*n* = 10), and 95% of redfish (*n* = 61 out of 64) caught on either the NE Greenland shelf or in the Fram Strait, were genetically assigned to the Barents Sea North East Arctic Cod (NEAC) population (Fig. 2c), and the Norwegian Shallow (NSH) redfish population (Fig. 2f), respectively. All shrimp (*n* = 40) caught on the NE Greenland shelf, were genetically assigned to the Spitsbergen West (SPW) shrimp population (Fig. 2i). Assignment with STRUCTURE was supported by high membership probabilities (*q* >0.8), which suggests that the three species on the NE Greenland shelf all originate from the Barents Sea. The only individuals to assign to an ‘Icelandic’ population were 5% of redfish (*n* = 3 out of 64).

There was high consistency in the identified population of origin between the assignment methods (STRUCTURE and *snapclust*), where 90% of cod, 98% of redfish and 75% of shrimp tests formed a consensus (Supplementary S5, Table S5). All individuals of these three species were assigned with a greater probability to the Barents Sea populations than any other reference population.

Discriminant Analysis of Principle Components (DAPC) strongly support the assignment results by clustering the NE Greenland specimens with the corresponding Barents Sea populations (Fig. 2b,e,h). The 95% DAPC cluster ellipses between NE Greenland and the Barents Sea overlapped considerably, though overlap was also evident between the reference populations, most significantly for the cod and shrimp clusters. The redfish and shrimp neighbour-joining trees resulted in the same grouping as the assignment and DAPC analyses, and indicate a Nei’s Distance of <0.02 between the redfish caught in NE Greenland and the Norwegian Shallow population (Fig. 3a). Nei’s Distance was as comparatively low (0.02) between the Norwegian and Icelandic shrimp reference populations as between the shrimp specimens of NE Greenland and the Spitsbergen West population (Fig. 3b).

**Fig. 3.**
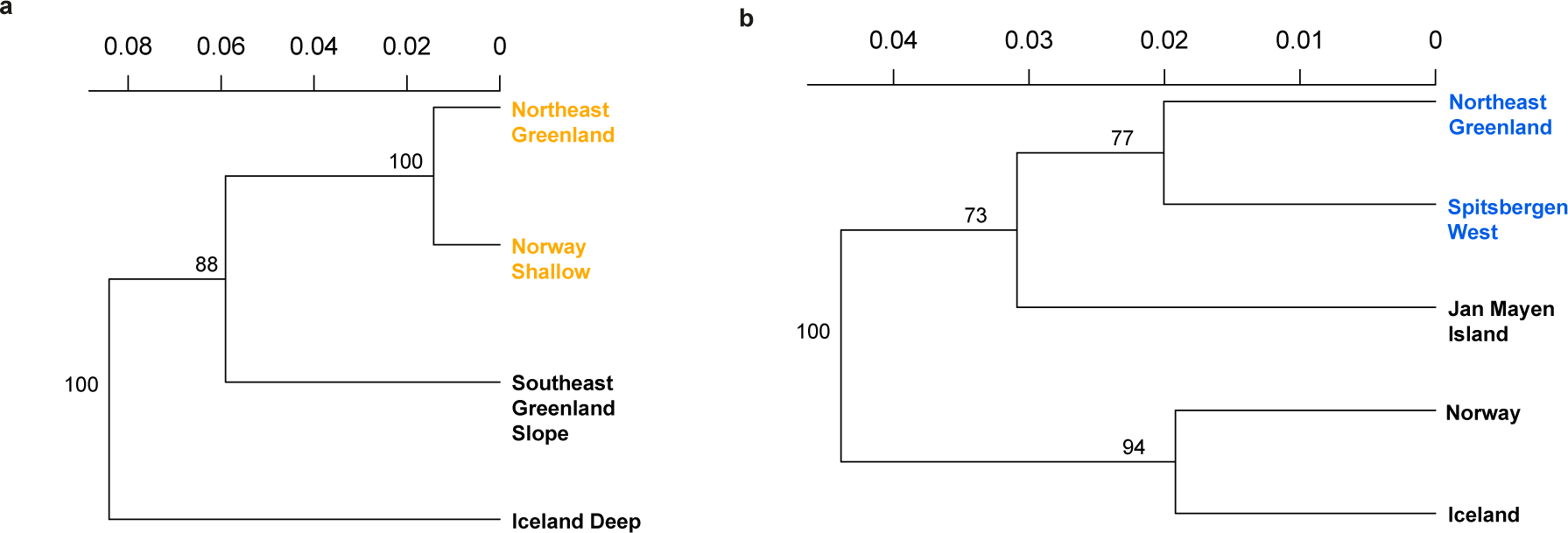
Neighbour-joining trees utilising Nei’s distance, for beaked redfish (*Sebastes mentella*) (**a**) and deep-sea shrimp (*Pandalus borealis*) (**b**) Northeast Greenland groups and reference populations. For abbreviations refer to Methods Table 2. Bootstrap values (>88% and >73%) on both trees suggest good reproducibility

## Discussion

Our results show that the NE Greenland shelf is readily reached by cod, redfish and shrimp from the Barents Sea, probably advected across the Fram Strait by the Return Atlantic Current, supporting recent simulation studies^25–27^. Advection plays an important role in the northward transport of plankton in the Barents Sea, via the West Spitsbergen Current^18^ and because up to 50% of this water is estimated to cross the Fram Strait^28–30^, the Return Atlantic Current provides a connection between the Barents Sea and the NE Greenland shelf ecosystems. The inflow of Atlantic water to the Barents Sea has doubled since 1980^31^, resulting in a warmer West Spitsbergen Current^32^. Hence, there is reason to believe that conditions on both sides of the Fram Strait have become more favourable for boreal species in recent years. The copepod *Calanus finmarchicus* is the major prey for young cod^33^ and its abundance during the last warm period in the North Atlantic (1920–1965) has likely driven the range expansion of cod and other boreal species^34^. Low abundances of *Calanus finmarchicus* were observed on the NE Greenland shelf in autumn 2006^35^, but in light of the West Spitsbergen Current warming, its abundance will likely increase in the Fram Strait and on the NE Greenland shelf^36^, thus providing ample food for boreal predators.

North East Arctic Cod (NEAC), the population of origin for the NE Greenland specimens, spawns along the Norwegian coast^37^ (latitudes 62–71 °N) during March and April where pelagic offspring drift by surface currents^38^ northwards and eastwards into the Barents Sea^39^. Depending on local wind-forcing, up to 1/3 of 0-group year-classes may advect off the Norwegian and Barents Sea shelf in some years and disperse over the Norwegian Sea^38^. We suggest those 0-group cod advected off the shelf by wind-forcing^27,40^ either outside of their spawning grounds, or at any point until their northern-most report west of Spitsbergen^21^, are particularly susceptible to cross the Fram Strait by the Return Atlantic Current (Fig. 2a). By October, when cod are >80 mm in total length (TL), they gain motility, descend out of the pelagic layer, and become demersal^42^. Therefore, for our explanation to hold true, 0-group cod from the Norwegian coast/Barents Sea must advect to the NE Greenland shelf by October of their spawning year. The observations of 0-group cod in the Fram Strait with a genetic signature of the NEAC population, in September 2007 and 2017, demonstrate that this is happening.

Redfish larvae are extruded along the continental shelf break of the Norwegian and Barents Seas from latitudes 64–74 °N between March and June^42^. Redfish larvae have been observed in Atlantic water west of the continental shelf, and as far north as Spitsbergen^23,43^. We observed large numbers of 0-group redfish (TL ∼40 mm) over the Fram Strait with a genetic signature of the Norwegian Shallow population. Juvenile redfish are pelagic until 40–50 mm TL at age 4–5 months when they gain motility and descend to deeper waters^44^. We propose that the 0-group redfish found in the Fram Strait in September 2017 were advected north to Spitsbergen along the shelf break by the West Spitsbergen Current, before crossing the Fram Strait by the Return Atlantic Current. The juvenile redfish had then reached the NE Greenland shelf along this route (Fig. 2d) by the time they were 4-5 months old.

Shrimp on the NE Greenland shelf also originated from the Barents Sea (see sampling of ^26^). Shrimp spawn in autumn throughout the Barents Sea and the meroplanktonic larvae hatch in spring. Shrimp larvae are highly-mobile, distributed according to currents until 2–3 months of age when they settle as post-larvae^14^. We find it more likely that shrimp larvae from the north-west Barents Sea, i.e. Spitsbergen, would reach the NE Greenland shelf than larvae from the northern Norwegian Coast or central-eastern Barents Sea, due to Spitsbergen’s close proximity to the Return Atlantic Current (Fig. 2g).

The NE Greenland shelf ecosystem is severely understudied and biodiversity baselines are fragmentary with no timeline^45^. It is therefore difficult to establish whether our findings reflect a recent shift driven by ocean warming or constitute a common component of the NE Greenland Shelf fauna. Nonetheless, the Barents Sea is the most productive ecosystem in the NE Atlantic^47^ and presently supports the historically largest cod population^41^. In addition, Atlantic herring (*Clupea harengus*), Atlantic haddock (*Melanogrammus aeglefinus*) and Atlantic mackerel (*Scomber scombrus*) are nowadays abundant in Spitsbergen waters. Therefore, in the future we could expect to find more boreal species on the NE Greenland shelf, exemplified by a recent observation of capelin (*Mallotus villosus*) in this area^13^. The three species studied herein are clearly not exceptional in being capable of entering the NE Greenland shelf. A recent simulation study^27^ demonstrates that between 2.4% and 12% of 0-group NEAC may be transported northwest along the proposed route (Fig. 2a). Advection has therefore the potential to restructure Arctic ecosystems^18^ and the route identified here suggests that a boreal faunal invasion of NE Greenland shelf from the Barents Sea is plausible. Trophic relationships are likely to be strongly modified^12^ as boreal generalists such as cod are favoured by climate scenarios^5^. Cod feed on polar cod (*Boreogadus saida*), Arctic seabed fishes and zoobenthos^11^, and as a figurehead of boreal range expansions into the Arctic, gives a glimpse of what is to come for native Arctic fauna.

This is the first report to disclose the genetic origin of boreal species in Arctic waters and the connection between Atlantic and Arctic ecosystems. Our findings support the hypothesis that cod, redfish, and shrimp disperse from the Barents Sea across the Fram Strait to the NE Greenland shelf. Due to a lack of time series, we cannot conclude if this is a new phenomenon, or not. In any case, predators and food competitors from lower latitudes alter trophic relationships and impact native Arctic fauna and, with a warming ocean in mind, we suggest that the NE Greenland shelf is likely to become populated by a larger proportion of boreal species.

## Methods

Specimens of juvenile cod (*Gadus morhua, n* = 7, body weight [bw]: 206–762 g), juvenile redfish (*Sebastes mentella, n* = 32, bw: 12–82 g), and adult shrimp (*Pandalus borealis, n* = 40) were caught via bottom trawl (c.f. ^14^) from 2007 to 2017 on the Northeast (NE) Greenland shelf (latitudes 74–79 °N), well outside of their known distributional range (Table 1). In addition, 0-group cod (*n* = 3) and 0-group redfish (*n* = 32) were caught via mid-water trawls (“Harstad” trawl, ∼20 min, ∼3 Knots) in the Fram Strait (Table 1) and are included in the analysis to support dispersal route hypotheses. Gill or muscle tissue samples from each specimen were preserved at sea in 96% ethanol and stored at −20 °C until further processing. Sampling was conducted using the R/V *Helmer Hanssen* as part of the TUNU-Programme^47^. A subset of (0-group) redfish and shrimp was used for genotyping, otherwise, genotyped individuals represent all specimens caught in the area. Access to sampling on the East Greenland shelf was permitted by the Government of Greenland under the remit of the TUNU-Programme at UiT, Norway.

**Table 1.** Details of assignment samples for each species. Station: p = Pelagic, b = Bottom, Year/Month = time of sampling. Totals for each species represent the number of genotyped individuals. Mean temp. = *in situ* sampling temperature obtained from CTD-sensor (Seabird 911).

| Station # | Latitude | Longitude | Year | Month | Mean temp. (°C) | Mean depth (m) | Cod | Redfish | Shrimp |
| --- | --- | --- | --- | --- | --- | --- | --- | --- | --- |
| 178 p | 76.55N | 03.03W | 2007 | 10 | 2.0 | 29 | 1 | - | - |
| 1312 b | 74.33N | 14.08W | 2015 | 8 | 1.8 | 300 | - | 11 | - |
| 1321 b | 75.09N | 13.38W | 2015 | 8 | 1.1 | 213 | 5 | 2 | - |
| 1339 b | 76.14N | 09.03W | 2015 | 8 | 1.6 | 280 | 1 | 7 | - |
| 1353 b | 77.28N | 05.49W | 2015 | 8 | 0.3 | 385 | 1 | 11 | 7 |
| 1278 p | 77.37N | 02.24E | 2017 | 9 | 5.6 | 34 | - | 16 | - |
| 1338 b | 76.00N | 14.18W | 2017 | 9 | 0.1 | 350 | - | 1 | 33 |
| 1381 p | 78.86N | 00.63W | 2017 | 9 | 1.2 | 26 | 2 | 16 | - |
| Genotyped individuals in total |  |  |  |  |  |  | 10 | 64 | 40 |

Genotyped reference populations (Table 2) for the Northeast Atlantic were obtained from several studies^16,26,48^. To ensure all relevant populations of each species in the Northeast Atlantic were well represented, the cod reference populations were supplemented by genotyping a representative cod population from Iceland, following the same procedure as listed below.

**Table 2.** Details of reference samples for each species. Year/Month = time of sampling. n = sample size (number of genotyped individuals).

| Species | Population | Year | Month | n | Latitude | Longitude |
| --- | --- | --- | --- | --- | --- | --- |
| Cod | Iceland | 2013 | 4 | 93 | 63.57N | 20.61W |
|  | Norwegian Coastal Cod | 2002 | 4 | 86 | 69.30N | 18.65E |
|  | North East Arctic Cod | 2005 | 12 | 47 | 74.10N | 21.10E |
|  | North East Arctic Cod | 2001 | 12 | 90 | 78.22N | 14.65E |
| Redfish | Iceland Deep | 2012 | 8 | 87 | 65.46N | 30.39W |
|  | South East Greenland Slope | 2011 | 3 | 133 | 64.24N | 35.14W |
|  | Norway Shallow | 2006 | 10 | 91 | 72.18N | 10.25E |
| Shrimp | Iceland | 2011 | 7 | 92 | 67.28N | 22.67W |
|  | Jan Mayen Island | 2011 | 10 | 88 | 70.61N | 08.43W |
|  | Norway | 2010 | 10 | 94 | 64.75N | 11.10E |
|  | Spitsbergen West | 2010 | 8 | 85 | 79.51N | 10.29E |

To our knowledge, the reference populations represent all the known spawning populations of these species within the NE Atlantic, which are relevant for this study. Cod and redfish are caught sporadically by commercial fishing in the waters around Jan Mayen Island. They are adult individuals fished during winter time (c.f. ^49^), which are suspected to originate from Icelandic and Barents Sea populations (winter feeding migration)^50^. Therefore, our reference populations should be adequate for assigning the juvenile cod and redfish considered in the present manuscript back to their population of origin.

DNA was isolated from ethanol-fixed gill or muscle tissue using the DNeasy Blood and Tissue Kit (Qiagen, Hilden, Germany) or the E-Z 96 Tissue DNA Kit (Omega Bio-Tek Inc., Norcross, GA, USA) following the manufacturer’s instructions.

Microsatellite loci were arranged in multiplexes (Supplementary Table S1), and amplified using polymerase chain reaction (PCR). PCR reactions (2.5 µL) contained ca. 1 x Qiagen Multiplex Master Mix, 0.1–1.0 µm primer, and 15–25 ng DNA. The 5’ end on the forward primers was labelled with a fluorescent dye by the manufacturer (Applied Biosystems, Foster City, CA, USA). Amplification was performed in a GeneAmp 2700 or 9700 thermal cycler (Applied Biosystems). PCR profiles were applied as per published protocols^16,48,51^ (Supplementary S1). PCR products were separated using an ABI 3130XL sequencer and GeneScan 500-LIZ (Applied Biosystems) was used as internal size standard. Alleles were automatically binned using GENEMAPPER software (v. 3.7, Applied Biosystems) and double-checked manually. Negative controls employed for extraction, amplification and fragmentation reported no contamination between samples. Replicates (33%) reported the repeatability and consistency of genotyping to be 100%.

Prior to analysis, reference genotypes that showed no amplification in >10% of loci were removed. This achieved amplification success >98% for each locus. All microsatellite loci were assessed for the presence of potential scoring errors, deviation from Hardy-Weinberg equilibrium (HWE), and non-neutrality (Supplementary S1). As the presence of scoring errors such as null alleles may introduce ambiguity around the true origin of the NE Greenland specimens, we ran analyses under two conditions, (1) removing loci showing potential scoring errors, and (2) inclusive of all loci (Supplementary S1, Table S1). This enabled us to retain loci subject to potential scoring errors where both conditions produced concurrent results, and to therefore minimise the loss of statistical power.

To increase the power of assignment (see Supplementary S2 for evaluation), only individuals with membership coefficients (*q*) lower/higher than 0.2/0.8 were used to establish reference population datasets (c.f. ^52^, Supplementary S3, Table S3). As weak population differentiation was expected within all datasets, we adopted a conservative approach to infer *q*^53^ using a no-admixture model as implemented in the Bayesian clustering method, STRUCTURE (v.2.3.4)^54^. This approach has been shown not to bias the true structuring in datasets with weak genetic differentiation^53^. STRUCTURE was run assuming no admixture (NOADMIX = 1), correlated allele frequencies (FREQSCORR = 1) and utilising locality data (LOCPRIOR = 1). The program was run using K = number of reference populations, for 10 iterations, each with a burn-in period and MCMC replicates of 500,000. CLUMPAK^55^ was used to merge runs (merged barplots: Supplementary S3, Fig. S3), and reported similarity scores >0.95.

STRUCTURE was employed as the principle tool to assign the NE Greenland individuals to previously identified populations. For this, STRUCTURE was run under the assignment mode (POPFLAG = 1), and assumed no admixture (NOADMIX = 1), correlated allele frequencies (FREQSCORR = 1) and utilised locality data (LOCPRIOR = 1). The program was run using K = number of reference populations, for 10 iterations, each with a burn-in period and MCMC replicates of 500,000. CLUMPAK reported run similarity scores >0.95. STRUCTURE barplots were visualised in R (v. 3.2.3)^56^ using the *pophelper* package (v. 2.2.5)^57^.

The maximum-likelihood clustering tool *snapclust*^58^, within the R package *adegenet* (v. 2.1.1)^59^, was used to corroborate the membership probabilities output by STRUCTURE. The function `snapclust` was run without optimization, and priors for the NE Greenland individuals were set to the reference population identified by STRUCTURE as the most probable origin. Runs used zero iterations (max.iter = 0) and membership coefficients were interpreted as output.

As an exploratory tool, Discriminant Analysis of Principle Components (DAPC)^60^, within the R package *adegenet*, was used to explore how the NE Greenland individuals relate to the reference populations. DAPC is a geometric clustering method free of HWE and linkage disequilibrium (LD) assumptions, that attempts to maximise the inter-variation between clusters while minimising the intra-variation observed within clusters.

DAPC clusters were set *a priori* to the number of reference populations plus one, including NE Greenland individuals as part of the DAPC model. The `x.val` function indicated the number of principle components (PC’s) to retain, but when this method resulted in the selection of too many PC’s, which would lead to overfitting, the `optim.a.score` function was preferred, based on an initial selection of all PC’s before refinement. All discriminant functions were retained due to the few clusters present (c.f. ^59^).

To identify the genetic distance between the NE Greenland individuals and reference populations, neighbour-joining trees were produced using the `aboot` function in the R package *poppr* (v. 2.3.0)^61^. This method utilised Nei’s Distance^62^ and 1000 bootstrap replicates. Due to the small sample size of NE Greenland cod, neighbour-joining trees were only produced for redfish (*n* = 64) and shrimp (*n* = 40) data.

Pre-analysis testing where loci subject to potential scoring errors were removed from analyses resulted in the same outcome as analysis retaining all loci (Supplementary S4). We therefore suggest that potential scoring errors had little impact on assignment and thus present our analyses utilising all loci available.

## Supporting information

## Acknowledgements

Thanks are extended to the Government of Greenland and the crew on board R/V *Helmer Hanssen* for access to the Greenland Sea. We thank Tanja L. Hanebrekke for laboratory assistance. This study is part of the TUNU-Programme, UiT The Arctic University of Norway, and was part-funded by a Small Research Grant from The Fisheries Society of the British Isles.

## Author Contributions

KP conceived the study. AJA, JSC, SB, AL, & CP collected tissues for analysis. AJA, JSC & KP designed the analysis. KP & JIW contributed data. AJA, SB & KP performed the analysis. AJA, JSC & KP wrote the manuscript. All authors reviewed the manuscript.

## Additional Information

None of the authors are subject to competing financial or non-financial interests in relation to the work.

