## Supplementary material for "Boreal marine fauna from the Barents Sea disperse to Arctic Northeast Greenland"

### Supplementary Text S1: PCR Profiles & Microsatellite loci evaluation

The PCR profile for cod (from [48]) consisted of an initial denaturation step at 95 °C for 15 min, followed by 22 cycles of 95 °C for 30 s, 56 °C for 3 min and 72 °C for 1 min. The PCR reactions ended with a final elongation step of 60 °C for 30 min.

The PCR profile for redfish (from [16]) consisted of an initial denaturation step at 95 °C for 15 min, followed by 25 cycles of 95 °C for 30 s, 56 °C for 90 s and 72 °C for 1 min. The PCR reactions ended with a final elongation step of 60 °C for 45 min.

The PCR profile for shrimp (from [51]) consisted of an initial denaturation step at 95 °C for 15 min, followed by 35 cycles of 95 °C for 30 s, 56 °C for 3 min and 72 °C for 1 min. The PCR reactions ended with a final elongation step of 60 °C for 30 min.

MICRO-CHECKER (v. 2.2.3)[63] was used to identify large allele drop-out, null alleles, and stuttering scoring errors. Locus-wise deviation from HWE was analysed using GENEPOP (v. 4.2.1)[64] using exact tests[65], in addition to Linkage Disequilibrium (LD) identification between loci. In addition, BAYESCAN (v. 2.1)[66] and ARLEQUIN (v. 3.5.2.2)[67] were used to test loci neutrality with default settings. ARLEQUIN simulations examined the joint distribution of $F_{ST}$ and heterozygosity under a hierarchical island model[68]. All results were judged for significance under the false discovery rate (FDR) approach[67] at the 5% level. Loci were not considered to be non-neutral unless both approaches reported them as such.

There was no evidence of large allele dropout in any loci. Several cases of stuttering and significant LD were reported, though these were deemed as non-genuine since genuine stuttering and LD are expected to affect all populations equally[70]. Loci were deemed to be subject to null alleles if null alleles were present in more than a single population. This applied to one locus of each species (cod: Gmo35, redfish: Spi6, shrimp: PbA104a). No loci deviated from HWE in more than a single population. No loci were consistently reported as non-neutral outliers under both approaches utilised.

**Table S1.** Microsatellite loci are shown in amplification multiplexes (MP's) and as utilised under both screening conditions and analyses (following: [16,48,51]).

| Species | Condition 1 | All loci |
|---|---|---|
| Cod | MP1: Gmo8, Gmo19, Gmo37, Tch11<br>MP2: Gmo2, Gmo3, Gmo34, Tch13, Gmo132 | MP1: Gmo8, Gmo19, Gmo35, Gmo37, Tch11<br>MP2: Gmo2, Gmo3, Gmo34, Tch13, Gmo132 |
| **Total** | **9** | **10** |
| Redfish | MP1: Sal1, Sal3, Sal4, Smen05<br>MP2: Spi4, Spi10, Smen10<br>MP3: Seb09, Seb25, Seb31, Seb33, Seb45 | MP1: Sal1, Sal3, Sal4, Smen05<br>MP2: Spi4, Spi6, Spi10, Smen10<br>MP3: Seb09, Seb25, Seb31, Seb33, Seb45 |
| **Total** | **12** | **13** |
| Shrimp | MP1: PbC8, PbC105, SD2-14<br>MP2: PbA1, PbA110, PbC109, PbD9<br>MP3: SD1-41, SD2-68, SD3-62 | MP1: PbA104a, PbC8, PbC105, SD2-14<br>MP2: PbA1, PbA110, PbC109, PbD9<br>MP3: SD1-41, SD2-68, SD3-62 |
| **Total** | **10** | **11** |

**Supplementary Text S2: Reference dataset evaluation**

To test the power of assignment, the function `predict.dapc` in the R package *adegenet* was used to re-assign all reference samples back to their original *a priori* population clusters. The `x.val` function indicated the number of principle components to retain. ARLEQUIN was used to calculate pairwise $F_{ST}$ values[71] between the reference populations. Only results for the all-loci condition of analysis are shown.

DAPC reassignment reported reference samples were successfully re-assigned to their original cluster (population) in 97% (cod) and 93% (redfish & shrimp) of cases. $F_{ST}$ values ranged from 0.011 to 0.040 and were all highly significant ($P$ <0.001) (Table S2).

**Table S2.** Pairwise $F_{ST}$ values and $P$-values. The values in boldface are significant after false discovery rate control at $P$ = 0.05. For abbreviations refer to Table S3.

| Species | | | | | |
|---|---|---|---|---|---|
| | | ICE | NCC | NEAC | |
| Cod | ICE | - | **0.000** | **0.000** | |
| | NCC | 0.024 | - | **0.000** | |
| | NEAC | 0.014 | 0.023 | - | |
| | | NSH | EGS | IDP | |
| Redfish | NSH | - | **0.000** | **0.000** | |
| | EGS | 0.031 | - | **0.000** | |
| | IDP | 0.037 | 0.040 | - | |
| | | NOR | SPW | ICE | JMA |
| Shrimp | NOR | - | **0.000** | **0.000** | **0.000** |
| | SPW | 0.031 | - | **0.000** | **0.000** |
| | ICE | 0.011 | 0.019 | - | **0.000** |
| | JMA | 0.038 | 0.024 | 0.025 | - |

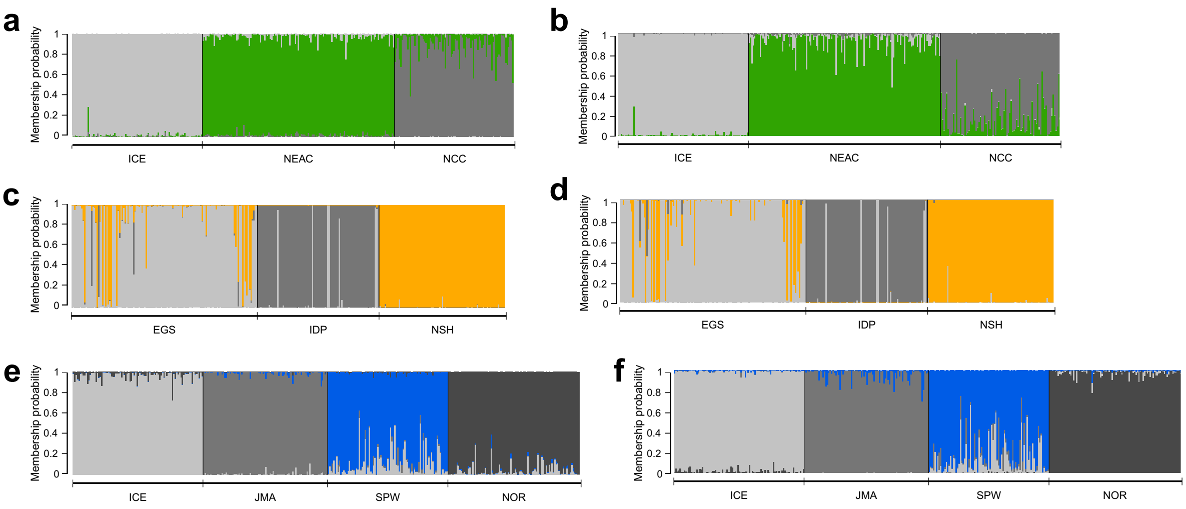

**Fig. S3.** STRUCTURE barplots showing Atlantic cod (*Gadus morhua*) (**a**, **b**), beaked redfish (*Sebastes mentella*) (**c**, **d**) and deep-sea shrimp (*Pandalus borealis*) (**e**, **f**) reference population membership probabilities (*q*) prior to the removal of individuals with a threshold of *q* lower/higher than 0.2/0.8. Barplots (**a**, **c** & **e**) utilised a reduced number of loci (Condition 1). Barplots (**b**, **d** & **f**) utilised all loci available regardless of potential scoring errors identified. For abbreviations refer to Table S3.

**Table S3.** Details of reference populations for each species, post removal of individuals with a threshold of *q* lower/higher than 0.2/0.8 and, under both analyses conditions. Abbr. = the abbreviated population name, *n* = sample size (number of genotyped individuals).

| Species | Population | Abbr. | Condition 1 *n* | All loci *n* |
|---|---|---|---|---|
| Cod | Iceland | ICE | 92 | 92 |
| | Norwegian Coastal Cod | NCC | 67 | 57 |
| | North East Arctic Cod | NEAC | 136 | 126 |
| Redfish | Iceland Deep | IDP | 80 | 80 |
| | South East Greenland Slope | EGS | 111 | 109 |
| | Norway Shallow | NSH | 91 | 90 |
| Shrimp | Iceland | ICE | 90 | 92 |
| | Jan Mayen Island | JMA | 87 | 87 |
| | Norway | NOR | 87 | 91 |
| | Spitsbergen West | SPW | 45 | 58 |

**Supplementary Text S4: Evaluation of the impact of potential scoring errors on assignment**

When using the datasets where loci subject to potential scoring errors (specifically, null alleles) were removed, individual assignment using STRUCTURE resulted in the same outcome as when analysing the data utilising all loci available, regardless of scoring errors. STRUCTURE barplots (Fig. S4) show that 100% of cod, 95% of redfish and 80% of shrimp tests resulted in the genetic assignment of individuals to a population within the Barents Sea, with a high membership probability ($q$ >0.8) suggesting a high-likelihood that specimens of the three species found off Northeast Greenland originate from the Barents Sea.

DAPC clustering using datasets where loci with potential scoring errors were removed resembled the results when using all loci available (Fig. S4a,c,e). The NE Greenland group of all three species clustered closely with the same Barents Sea populations as indicated by the assignment testing (Fig. S4b,d,f). The 95% DAPC cluster ellipses between NEG and these population clusters overlapped considerably, though overlap was also evident between the reference population clusters, most significantly for the cod and shrimp clusters. The distance between the NE Greenland group clusters and clusters they were assigned to using STRUCTURE only differed for the shrimp data. Here, there is a greater distance between the NE Greenland shrimp group and Spitsbergen West shrimp group, than when analysed using all loci available.

Therefore, we suggest that the impact of potential null alleles on assignment was minimal and that utilising all loci available regardless of null alleles did not bias the assignment outcome.

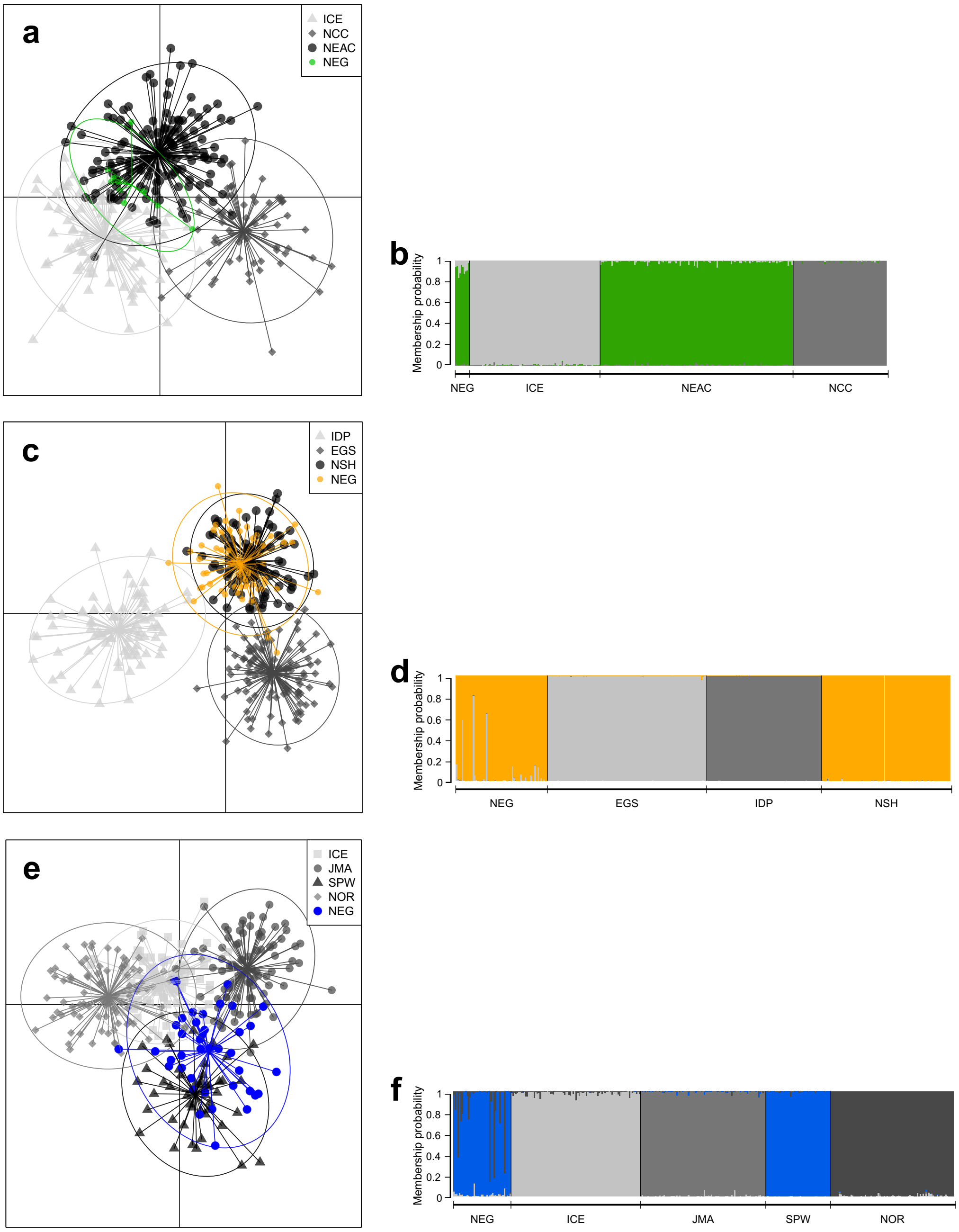

**Fig. S4.** Genetic analysis using a reduced set of loci that accounted for potential scoring errors, that we termed 'Condition 1', for Atlantic cod (*Gadus morhua*) (**a**, **b**), beaked redfish (*Sebastes mentella*) (**c**, **d**) and deep-sea shrimp (*Pandalus borealis*) (**e**, **f**) data. DAPC scatterplots (**a, c**, **e**) show how the NE Greenland groups relate to the reference populations of the Northeast Atlantic Ocean. DAPC cluster ellipses were set to contain 95% of genotypes. DAPC scatterplots explain 91% (**a**), 94% (**c**) and 99% (**e**) of the total variation observed. STRUCTURE barplots (**b**, **d**, **f**) show membership probabilities (*q*) for NE Greenland individuals based on the reference populations used. For abbreviations refer to Table S3.

### Supplementary Text S5: Assignment membership probabilities

STRUCTURE and *snapclust* membership probabilities ($q$) for cod (Table S5.1), redfish (Table S5.2) and shrimp (Table S5.3) specimens caught off Northeast Greenland are shown, and summarised with the most probable assignment origin indicated for both methods.

**Table S5.1.** STRUCTURE and *snapclust* identified cod population of origin and membership probabilities ($q$) for Northeast Greenland (NEG) cod individuals using all loci. Bold typeface indicates the $q$ values supporting the most probable origin of each individual. For abbreviations refer to Table S3.

| Sample | STRUCTURE origin | *snapclust* origin | STRUCTURE $q$ values | | | *snapclust* $q$ values | | |
|---|---|---|---|---|---|---|---|---|
| | | | ICE | NEAC | NCC | ICE | NEAC | NCC |
| NEG Cod 1 | NEAC | NEAC | 0.133 | **0.852** | 0.015 | 0.230 | **0.630** | 0.140 |
| NEG Cod 2 | NEAC | NEAC | 0.074 | **0.925** | 0.001 | 0.056 | **0.943** | 0.001 |
| NEG Cod 3 | NEAC | NEAC | 0.158 | **0.839** | 0.003 | 0.388 | **0.601** | 0.011 |
| NEG Cod 4 | NEAC | NEAC | 0.148 | **0.848** | 0.003 | 0.380 | **0.605** | 0.015 |
| NEG Cod 5 | NEAC | NEAC | 0.063 | **0.935** | 0.002 | 0.092 | **0.887** | 0.021 |
| NEG Cod 6 | NEAC | NEAC | 0.160 | **0.835** | 0.005 | 0.373 | **0.620** | 0.007 |
| NEG Cod 7 | NEAC | NEAC | 0.144 | **0.840** | 0.016 | 0.284 | **0.631** | 0.085 |
| NEG Cod 8 | NEAC | NEAC | 0.171 | **0.825** | 0.004 | 0.466 | **0.520** | 0.014 |
| NEG Cod 9 | NEAC | ICE | 0.188 | **0.802** | 0.010 | **0.557** | 0.326 | 0.117 |
| NEG Cod 10 | NEAC | NEAC | 0.038 | **0.962** | 0.000 | 0.036 | **0.962** | 0.002 |

**Table S5.2.** STRUCTURE and *snapclust* identified redfish population of origin and membership probabilities (*q*) for Northeast Greenland (NEG) redfish individuals using all loci. Bold typeface indicates the *q* values supporting the most probable origin of each individual. For abbreviations refer to Table S3.

| Sample | STRUCTURE origin | *snapclust* origin | STRUCTURE *q* values | | | *snapclust q* values | | |
|---|---|---|---|---|---|---|---|---|
| | | | EGS | IDP | NSH | EGS | IDP | NSH |
| NEG Redfish 1 | NSH | NSH | 0.044 | 0.000 | **0.956** | 0.217 | 0.001 | **0.782** |
| NEG Redfish 2 | NSH | NSH | 0.000 | 0.000 | **1.000** | 0.001 | 0.016 | **0.983** |
| NEG Redfish 3 | NSH | NSH | 0.003 | 0.002 | **0.995** | 0.036 | 0.056 | **0.908** |
| NEG Redfish 4 | NSH | NSH | 0.001 | 0.000 | **0.999** | 0.004 | 0.001 | **0.995** |
| NEG Redfish 5 | EGS | EGS | **0.870** | 0.052 | 0.079 | **0.553** | 0.226 | 0.221 |
| NEG Redfish 6 | NSH | NSH | 0.004 | 0.000 | **0.996** | 0.044 | 0.000 | **0.956** |
| NEG Redfish 7 | NSH | NSH | 0.000 | 0.000 | **1.000** | 0.001 | 0.000 | **0.999** |
| NEG Redfish 8 | NSH | NSH | 0.000 | 0.000 | **1.000** | 0.001 | 0.000 | **0.999** |
| NEG Redfish 9 | NSH | NSH | 0.000 | 0.000 | **1.000** | 0.001 | 0.000 | **0.999** |
| NEG Redfish 10 | NSH | NSH | 0.000 | 0.000 | **1.000** | 0.001 | 0.000 | **0.999** |
| NEG Redfish 11 | NSH | NSH | 0.000 | 0.000 | **1.000** | 0.000 | 0.000 | **1.000** |
| NEG Redfish 12 | NSH | NSH | 0.000 | 0.000 | **1.000** | 0.012 | 0.001 | **0.987** |
| NEG Redfish 13 | EGS | EGS | **0.919** | 0.000 | 0.081 | **0.990** | 0.000 | 0.010 |
| NEG Redfish 14 | NSH | NSH | 0.027 | 0.002 | **0.971** | 0.017 | 0.032 | **0.801** |
| NEG Redfish 15 | NSH | NSH | 0.003 | 0.000 | **0.997** | 0.023 | 0.000 | **0.977** |
| NEG Redfish 16 | NSH | NSH | 0.003 | 0.001 | **0.996** | 0.031 | 0.061 | **0.908** |
| NEG Redfish 17 | NSH | NSH | 0.000 | 0.000 | **1.000** | 0.001 | 0.000 | **0.999** |
| NEG Redfish 18 | NSH | NSH | 0.000 | 0.000 | **1.000** | 0.001 | 0.001 | **0.998** |
| NEG Redfish 19 | NSH | NSH | 0.001 | 0.000 | **0.999** | 0.002 | 0.000 | **0.998** |
| NEG Redfish 20 | NSH | NSH | 0.005 | 0.000 | **0.995** | 0.063 | 0.000 | **0.937** |
| NEG Redfish 21 | NSH | NSH | 0.000 | 0.000 | **1.000** | 0.009 | 0.000 | **0.991** |
| NEG Redfish 22 | EGS | EGS | **0.746** | 0.001 | 0.253 | **0.953** | 0.002 | 0.045 |
| NEG Redfish 23 | NSH | NSH | 0.001 | 0.000 | **0.999** | 0.023 | 0.003 | **0.974** |
| NEG Redfish 24 | NSH | NSH | 0.001 | 0.000 | **0.999** | 0.033 | 0.003 | **0.964** |
| NEG Redfish 25 | NSH | NSH | 0.000 | 0.000 | **1.000** | 0.001 | 0.000 | **0.999** |
| NEG Redfish 26 | NSH | NSH | 0.000 | 0.000 | **1.000** | 0.009 | 0.000 | **0.991** |
| NEG Redfish 27 | NSH | NSH | 0.000 | 0.000 | **1.000** | 0.001 | 0.000 | **0.999** |
| NEG Redfish 28 | NSH | NSH | 0.006 | 0.000 | **0.994** | 0.048 | 0.000 | **0.952** |
| NEG Redfish 29 | NSH | NSH | 0.000 | 0.000 | **1.000** | 0.005 | 0.000 | **0.995** |
| NEG Redfish 30 | NSH | NSH | 0.000 | 0.000 | **1.000** | 0.001 | 0.001 | **0.998** |
| NEG Redfish 31 | NSH | NSH | 0.000 | 0.000 | **1.000** | 0.007 | 0.000 | **0.993** |

| | | | | | | | | |
|---|---|---|---|---|---|---|---|---|
| NEG Redfish 32 | NSH | NSH | 0.000 | 0.000 | **1.000** | 0.002 | 0.000 | **0.998** |
| NEG Redfish 33 | NSH | NSH | 0.000 | 0.000 | **1.000** | 0.000 | 0.000 | **0.999** |
| NEG Redfish 34 | NSH | NSH | 0.008 | 0.000 | **0.992** | 0.076 | 0.000 | **0.924** |
| NEG Redfish 35 | NSH | NSH | 0.009 | 0.001 | **0.990** | 0.028 | 0.024 | **0.948** |
| NEG Redfish 36 | NSH | NSH | 0.000 | 0.000 | **1.000** | 0.004 | 0.000 | **0.996** |
| NEG Redfish 37 | NSH | NSH | 0.000 | 0.000 | **1.000** | 0.026 | 0.003 | **0.971** |
| NEG Redfish 38 | NSH | NSH | 0.000 | 0.000 | **1.000** | 0.002 | 0.000 | **0.998** |
| NEG Redfish 39 | NSH | NSH | 0.000 | 0.000 | **1.000** | 0.001 | 0.000 | **0.999** |
| NEG Redfish 40 | NSH | EGS | 0.120 | 0.000 | **0.880** | **0.693** | 0.000 | 0.307 |
| NEG Redfish 41 | NSH | NSH | 0.022 | 0.000 | **0.978** | 0.023 | 0.000 | **0.977** |
| NEG Redfish 42 | NSH | NSH | 0.004 | 0.000 | **0.996** | 0.176 | 0.005 | **0.819** |
| NEG Redfish 43 | NSH | NSH | 0.000 | 0.000 | **1.000** | 0.000 | 0.000 | **1.000** |
| NEG Redfish 44 | NSH | NSH | 0.000 | 0.000 | **1.000** | 0.000 | 0.001 | **0.999** |
| NEG Redfish 45 | NSH | NSH | 0.000 | 0.000 | **1.000** | 0.001 | 0.000 | **0.999** |
| NEG Redfish 46 | NSH | NSH | 0.028 | 0.000 | **0.971** | 0.127 | 0.002 | **0.871** |
| NEG Redfish 47 | NSH | NSH | 0.000 | 0.000 | **1.000** | 0.012 | 0.000 | **0.988** |
| NEG Redfish 48 | NSH | NSH | 0.001 | 0.002 | **0.997** | 0.005 | 0.045 | **0.950** |
| NEG Redfish 49 | NSH | NSH | 0.001 | 0.000 | **0.999** | 0.007 | 0.000 | **0.993** |
| NEG Redfish 50 | NSH | NSH | 0.000 | 0.000 | **1.000** | 0.000 | 0.001 | **0.999** |
| NEG Redfish 51 | NSH | NSH | 0.000 | 0.000 | **1.000** | 0.000 | 0.001 | **0.999** |
| NEG Redfish 52 | NSH | NSH | 0.000 | 0.000 | **1.000** | 0.001 | 0.000 | **0.999** |
| NEG Redfish 53 | NSH | NSH | 0.004 | 0.000 | **0.996** | 0.175 | 0.000 | **0.825** |
| NEG Redfish 54 | NSH | NSH | 0.000 | 0.000 | **1.000** | 0.000 | 0.000 | **1.000** |
| NEG Redfish 55 | NSH | NSH | 0.000 | 0.000 | **1.000** | 0.000 | 0.000 | **1.000** |
| NEG Redfish 56 | NSH | NSH | 0.000 | 0.000 | **1.000** | 0.285 | 0.000 | **0.715** |
| NEG Redfish 57 | NSH | NSH | 0.000 | 0.000 | **1.000** | 0.000 | 0.000 | **1.000** |
| NEG Redfish 58 | NSH | NSH | 0.002 | 0.000 | **0.998** | 0.107 | 0.001 | **0.892** |
| NEG Redfish 59 | NSH | NSH | 0.000 | 0.000 | **1.000** | 0.000 | 0.001 | **0.999** |
| NEG Redfish 60 | NSH | NSH | 0.001 | 0.000 | **0.999** | 0.025 | 0.000 | **0.975** |
| NEG Redfish 61 | NSH | NSH | 0.000 | 0.000 | **1.000** | 0.000 | 0.004 | **0.996** |
| NEG Redfish 62 | NSH | NSH | 0.000 | 0.000 | **1.000** | 0.016 | 0.010 | **0.974** |
| NEG Redfish 63 | NSH | NSH | 0.000 | 0.000 | **1.000** | 0.007 | 0.000 | **0.993** |
| NEG Redfish 64 | NSH | NSH | 0.000 | 0.000 | **1.000** | 0.000 | 0.000 | **1.000** |

**Table S5.3.** STRUCTURE and *snapclust* identified shrimp population of origin and membership probabilities ($q$) for Northeast Greenland (NEG) shrimp individuals using all loci. Bold typeface indicates the $q$ values supporting the most probable origin of each individual. For abbreviations refer to Table S3.

| Sample | STRUCTURE origin | *snapclust* origin | STRUCTURE $q$ values | | | | *snapclust* $q$ values | | | |
|---|---|---|---|---|---|---|---|---|---|---|
| | | | ICE | JMA | SPW | NOR | ICE | JMA | SPW | NOR |
| NEG Shrimp 1 | SPW | ICE | 0.027 | 0.027 | **0.938** | 0.008 | **0.451** | 0.439 | 0.096 | 0.014 |
| NEG Shrimp 2 | SPW | SPW | 0.008 | 0.000 | **0.976** | 0.016 | 0.063 | 0.012 | **0.751** | 0.175 |
| NEG Shrimp 3 | SPW | SPW | 0.008 | 0.015 | **0.977** | 0.000 | 0.301 | 0.253 | **0.445** | 0.002 |
| NEG Shrimp 4 | SPW | NOR | 0.003 | 0.02 | **0.914** | 0.063 | 0.010 | 0.168 | 0.207 | **0.615** |
| NEG Shrimp 5 | SPW | SPW | 0.015 | 0.001 | **0.982** | 0.002 | 0.197 | 0.022 | **0.772** | 0.009 |
| NEG Shrimp 6 | SPW | SPW | 0.007 | 0.008 | **0.962** | 0.023 | 0.135 | 0.132 | **0.534** | 0.200 |
| NEG Shrimp 7 | SPW | NOR | 0.006 | 0.000 | **0.979** | 0.015 | 0.159 | 0.009 | 0.257 | **0.575** |
| NEG Shrimp 8 | SPW | SPW | 0.011 | 0.001 | **0.983** | 0.005 | 0.169 | 0.018 | **0.715** | 0.098 |
| NEG Shrimp 9 | SPW | SPW | 0.004 | 0.000 | **0.996** | 0.000 | 0.028 | 0.012 | **0.958** | 0.001 |
| NEG Shrimp 10 | SPW | SPW | 0.007 | 0.000 | **0.987** | 0.006 | 0.137 | 0.001 | **0.805** | 0.057 |
| NEG Shrimp 11 | SPW | SPW | 0.002 | 0.013 | **0.967** | 0.018 | 0.008 | 0.349 | **0.584** | 0.058 |
| NEG Shrimp 12 | SPW | SPW | 0.000 | 0.000 | **0.998** | 0.002 | 0.001 | 0.003 | **0.967** | 0.028 |
| NEG Shrimp 13 | SPW | SPW | 0.000 | 0.001 | **0.999** | 0.000 | 0.000 | 0.098 | **0.900** | 0.001 |
| NEG Shrimp 14 | SPW | SPW | 0.001 | 0.000 | **0.998** | 0.001 | 0.002 | 0.002 | **0.991** | 0.005 |
| NEG Shrimp 15 | SPW | SPW | 0.005 | 0.001 | **0.990** | 0.004 | 0.023 | 0.026 | **0.832** | 0.119 |
| NEG Shrimp 16 | SPW | ICE | 0.042 | 0.003 | **0.954** | 0.001 | **0.798** | 0.013 | 0.188 | 0.002 |
| NEG Shrimp 17 | SPW | JMA | 0.000 | 0.042 | **0.958** | 0.000 | 0.001 | **0.707** | 0.290 | 0.002 |
| NEG Shrimp 18 | SPW | SPW | 0.001 | 0.014 | **0.985** | 0.000 | 0.003 | 0.273 | **0.716** | 0.007 |
| NEG Shrimp 19 | SPW | SPW | 0.002 | 0.001 | **0.997** | 0.000 | 0.042 | 0.018 | **0.933** | 0.007 |
| NEG Shrimp 20 | SPW | SPW | 0.001 | 0.007 | **0.992** | 0.000 | 0.008 | 0.437 | **0.555** | 0.000 |
| NEG Shrimp 21 | SPW | SPW | 0.008 | 0.004 | **0.986** | 0.001 | 0.208 | 0.171 | **0.551** | 0.070 |
| NEG Shrimp 22 | SPW | SPW | 0.007 | 0.001 | **0.992** | 0.000 | 0.180 | 0.055 | **0.751** | 0.014 |
| NEG Shrimp 23 | SPW | SPW | 0.001 | 0.001 | **0.999** | 0.000 | 0.007 | 0.033 | **0.960** | 0.001 |
| NEG Shrimp 24 | SPW | SPW | 0.000 | 0.000 | **1.000** | 0.000 | 0.000 | 0.028 | **0.971** | 0.000 |
| NEG Shrimp 25 | SPW | SPW | 0.004 | 0.002 | **0.993** | 0.001 | 0.052 | 0.095 | **0.846** | 0.007 |
| NEG Shrimp 26 | SPW | NOR | 0.013 | 0.001 | **0.824** | 0.162 | 0.017 | 0.004 | 0.038 | **0.942** |
| NEG Shrimp 27 | SPW | SPW | 0.004 | 0.008 | **0.979** | 0.009 | 0.041 | 0.126 | **0.776** | 0.057 |
| NEG Shrimp 28 | SPW | SPW | 0.007 | 0.002 | **0.990** | 0.001 | 0.112 | 0.050 | **0.828** | 0.010 |
| NEG Shrimp 29 | SPW | NOR | 0.004 | 0.000 | **0.924** | 0.072 | 0.022 | 0.001 | 0.212 | **0.765** |
| NEG Shrimp 30 | SPW | SPW | 0.005 | 0.003 | **0.976** | 0.016 | 0.079 | 0.037 | **0.751** | 0.132 |
| NEG Shrimp 31 | SPW | SPW | 0.003 | 0.005 | **0.991** | 0.001 | 0.021 | 0.201 | **0.769** | 0.009 |

| | | | | | | | | | | |
|---|---|---|---|---|---|---|---|---|---|---|
| NEG Shrimp 32 | SPW | SPW | 0.011 | 0.003 | **0.985** | 0.001 | 0.203 | 0.144 | **0.645** | 0.008 |
| NEG Shrimp 33 | SPW | JMA | 0.005 | 0.026 | **0.945** | 0.024 | 0.029 | **0.587** | 0.273 | 0.111 |
| NEG Shrimp 34 | SPW | SPW | 0.003 | 0.000 | **0.996** | 0.001 | 0.183 | 0.004 | **0.798** | 0.016 |
| NEG Shrimp 35 | SPW | SPW | 0.004 | 0.000 | **0.996** | 0.000 | 0.128 | 0.002 | **0.869** | 0.002 |
| NEG Shrimp 36 | SPW | ICE | 0.052 | 0.008 | **0.837** | 0.103 | **0.504** | 0.016 | 0.069 | 0.411 |
| NEG Shrimp 37 | SPW | SPW | 0.000 | 0.000 | **1.000** | 0.000 | 0.000 | 0.001 | **0.999** | 0.000 |
| NEG Shrimp 38 | SPW | SPW | 0.004 | 0.003 | **0.993** | 0.000 | 0.013 | 0.238 | **0.749** | 0.000 |
| NEG Shrimp 39 | SPW | SPW | 0.002 | 0.002 | **0.996** | 0.000 | 0.030 | 0.109 | **0.851** | 0.010 |
| NEG Shrimp 40 | SPW | JMA | 0.008 | 0.054 | **0.937** | 0.001 | 0.094 | **0.680** | 0.225 | 0.002 |
